# Evidence for long-standing mild-effect mutator alleles shaping genetic variation in *S. cerevisiae* populations

**DOI:** 10.1101/2025.11.22.689970

**Authors:** Pengyao Jiang, Vidha Sudhesh, Megan My-Ngan Phan, Ishan Bansal, Cian Wheeler, Alan J Herr, Maitreya J. Dunham, Kelley Harris

## Abstract

Most mutations are neutral or deleterious, and mutator alleles that substantially increase the mutation rate of an organism are believed to be short-lived in natural populations. Theory suggests that mutator alleles of modest effect have the potential to segregate nearly neutrally, but this prediction has been difficult to test experimentally. Here, we report strong genomic signatures consistent with the transmission and long-term maintenance of at least one natural mutator allele in *Saccharomyces cerevisiae* isolates. Specifically, such signatures are consistent with segregation of mutator alleles that disproportionately increase A>C mutations in the African Beer strains of *S. cerevisiae*. Remarkably, the mutation spectra of African Beer strains deviate from that of other *S. cerevisiae* natural isolates more than some *Saccharomyces* species differ from each other. Furthermore, computational analysis suggests mutator allele introgression from the African Beer population into a subset of French Dairy yeast, motivating us to experimentally measure the *de novo* mutation rates and spectra of these strains. We observed a consistent but weak enrichment of A>C and A>G *de novo* mutations in the African Beer population, with the A>C enrichment also evident in the French Dairy subset. Unexpectedly, other *de novo* mutation types varied more substantially among strains, including one outlier African Beer strain (AFL) with an excess of C>A mutations. Overall, our measurements reveal that *de novo* mutation spectra show greater variability among populations compared to the mutation spectra of rare polymorphisms, which in turn show greater variability than the spectra of all polymorphisms. Together, while additional mutator alleles may segregate in natural populations, mild enrichment of A>C and A>G types in African Beer strains reflects mutator alleles with weak effects that can persist and accumulate mutations through evolution.

## Introduction

Mutations are the ultimate source of genetic variation and evolution (Darwin 1859; Simpson 1944; Muller 1950), but most of them are neutral or deleterious. To minimize the harmful consequences of mutations, organisms have evolved complex molecular mechanisms that maintain extremely low mutation rates, ensuring the faithful transmission of genetic material across generations (Beckman and Loeb 1993; Eisen and Hanawalt 1999). Historically, population geneticists have treated the mutation rate as a constant (Kimura 1983; Gillespie 2004; Lawson et al. 2012), an approximation that enabled the development of powerful inference methods based on the principle of the molecular clock (Gutenkunst et al. 2009; Sprengelmeyer et al. 2020; Wilson and Sarich 1969). However, the existence of mutation rate variation is well established (Sturtevant 1937; Li et al. 1987), and mutation rates vary by nearly four orders of magnitude across the tree of life (Lynch et al. 2023; Lynch 2010), indicating that the mutation rate itself evolves (Melde et al. 2022). The drift-barrier hypothesis has been proposed to explain the equilibrium mutation rate across species (Lynch et al. 2023; Lynch 2010). However, despite this progress, the evolutionary dynamics of mutation rate variation and evolution in natural populations remain poorly understood.

Hypermutator strains frequently arise during the experimental evolution of microbial populations (Sniegowski et al. 1997; Blount et al. 2018) and in natural populations of bacterial (Pal et al. 2007) and fungal pathogens (Billmyre et al. 2017; Boyce et al. 2017; Healey et al. 2016), which can easily increase the mutation rate by 10 to 100 fold (Pal et al. 2007). Similar increases in somatic mutation rates are seen in human cancer, but they vary to a large extent depending on the cancer type (Watson et al. 2013; Milholland et al. 2017). Such mutators may provide a selective advantage in novel, stressful, or fluctuating environments (Ram and Hadany 2012; Steenwyk et al. 2019). Nevertheless, because of the fitness costs associated with an increased mutation load, mutator alleles are generally short-lived, as observed in experimental evolution studies (McDonald et al. 2012).

In free-living non-pathogenic organisms, population-level variation in heritable mutation rate and spectrum is generally modest (Bergeron et al. 2023; Wang and Obbard 2023). In humans, for example, an enrichment of the TCC>TTC triplet mutation in European populations is the relic of a mutator phenotype that is no longer active and was likely associated with an overall mutation rate change of less than 1.1-fold (Harris and Pritchard 2017). Only a handful of studies have characterized natural mutator alleles, and these mutator alleles are either extremely rare or have a small effect size (Jiang et al. 2021; Sasani et al. 2022, 2024; Kaplanis et al. 2022). In *Saccharomyces cerevisiae*, reporter gene assays suggest that the mutation rates of natural isolates vary by tenfold, and some of this variation is driven by a natural mutator affecting C>A mutations (Jiang et al. 2021)). Similarly, in mice, at least two natural mutator alleles affect the rate of C>A mutations, but again together these effects cause less than a 10% increase in overall mutation rate (Sasani et al. 2022, 2024).

Patterns in mutation spectra inferred from natural polymorphisms can sometimes provide insights into population history. Admixture is one potential explanation for intermediate positions on mutational PCA plots, as the mixing of genetically differentiated populations can generate composite mutation spectra. For example, in humans, the African American and admixed Amerindian populations exhibit mutation spectra resembling mixtures of European, African, and East Asian spectra (Harris and Pritchard 2017). Consequently, methods for inferring local ancestry resulting from admixture (Lawson et al. 2012) can be used alongside mutation- spectrum analyses to reconstruct historical changes in mutational processes. Recently, a genealogy-based framework was developed that jointly infers local ancestry and admixture from unsampled ("ghost") populations (Loya et al. 2026). By integrating ancestry-specific mutational spectra with genome-wide genealogies, the method identifies the origins and movements of admixture tracts across Africa and Eurasia and reveals multiple deep admixture events in human history.

Here, we report a novel pattern indicating the persistence and transmission of natural mutator alleles in *S. cerevisiae* isolates. We identify these patterns by studying mutation spectrum variation in the 1,011 *S. cerevisiae* collection, a public resource of *S. cerevisiae* strains that were collected worldwide from diverse natural and domesticated niches (Peter et al. 2018). We previously reported that the relative rates of different base substitution types exhibited variation across this dataset that reflected the population structure of *S. cerevisiae*. In addition, we found that strains from African Beer had a highly unusual spectrum that stood out from this background variation, with disproportionately high levels of A>C and A>G mutations (Jiang et al. 2021). In this study, we show that this signature is also present in a subset of the French Dairy strains, a pattern that is consistent with the introgression of one or more mutators from African Beer into the French Dairy population. We verify that these mutator effects are measurable in *de novo* mutations accumulated in laboratory isolates of these strains, though the effects of these alleles appear extremely weak. These results show that natural variation in African Beer of *S. cerevisiae* is likely shaped by weak, long-lived mutator alleles, indicating that natural selection has limited power to fully optimize mutation rates within a species, even over evolutionary timescales.

## Results

### The *S. cerevisiae* African Beer population has an unusual spectrum of segregating mutations

To investigate mutation spectrum variation among natural yeast isolates, we previously performed principal component analysis (PCA) on mutation spectra derived from natural polymorphisms across 1,011 *S. cerevisiae* isolates (Figure 1A of (Jiang et al. 2021)). We found that the African Beer population exhibits an outlier mutation spectrum compared with other populations, showing enrichment for A>G and A>C mutations along PC1 and PC2, respectively. A subset of the French Dairy strains also exhibited intermediate enrichment towards the African Beer direction.

**Figure 1.**
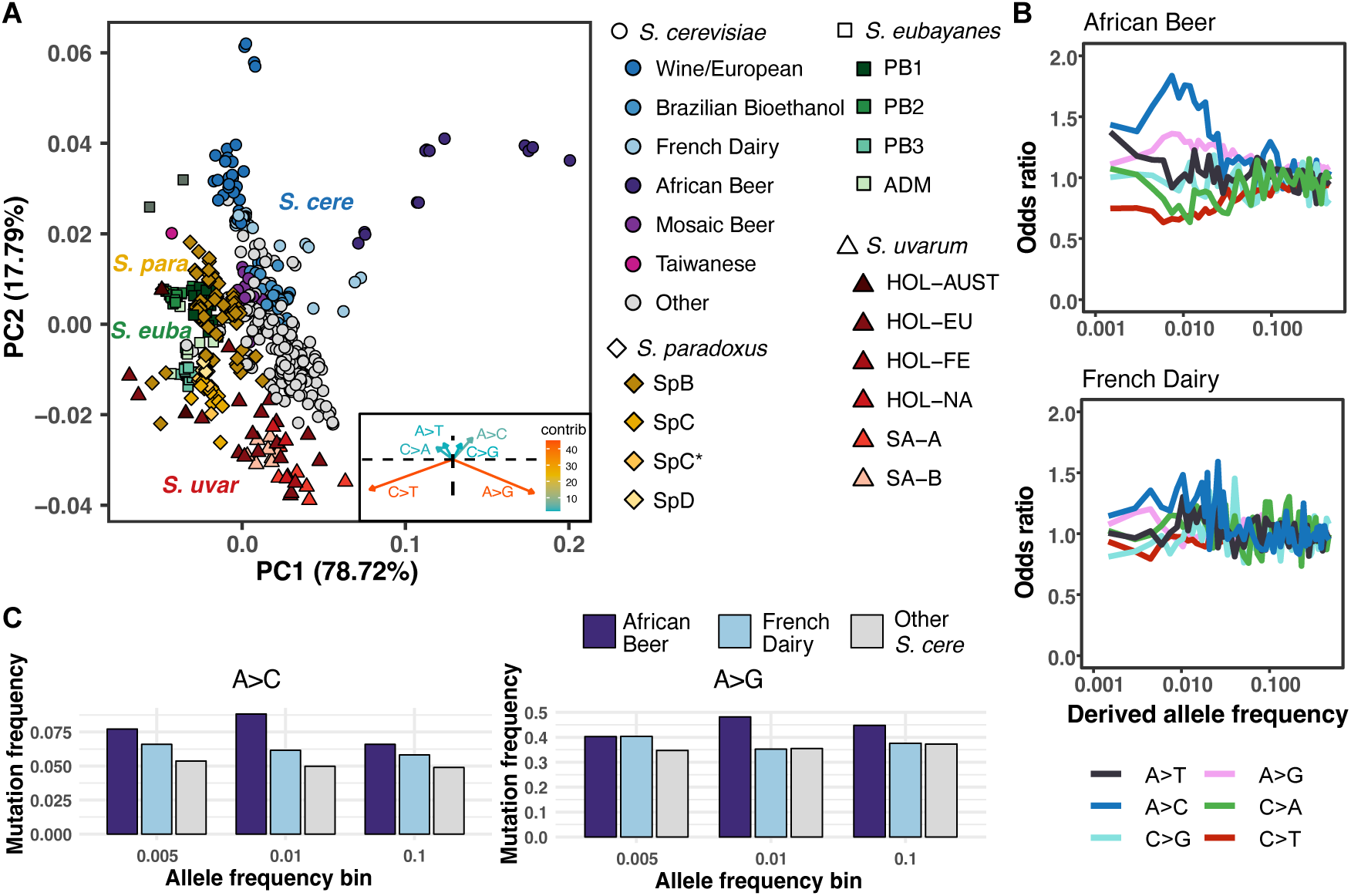
***S. cerevisiae* African Beer population exhibits an extreme outlier mutation spectrum compared with other *Saccharomyces* yeasts.** (A) Mutational PCA from 4 species in the *Saccharomyces* genus, including *S. cerevisiae, S. paradoxus, S. eubayanus,* and *S. uvarum*. Each strain is shown as a single dot, colored by previously defined populations. *S. cerevisiae* strains are shown as circles, *S. paradoxus* strains as diamonds, *S. eubayanus* as squares, and *S. uvarum* as triangles. Inset: The loadings of this mutation spectrum PCA show that the African Beer strains are enriched for A>G and A>C mutations. (B) Odds ratio of the mutation frequency of African Beer and French Dairy, by dividing the mutation frequency relative to the average of all other *S. cerevisiae* among each derived allele frequency bin. (C) A>C and A>G mutation frequency differences in African Beer, French Dairy, and all other *S. cerevisiae* across allele frequency bins.

To contextualize the magnitude of mutation spectrum divergence between the African Beer group and other *S. cerevisiae* strains, we assayed mutation spectrum variation across species in the genus *Saccharomyces*. We analyzed single-nucleotide polymorphisms (SNPs) from species represented by published population datasets of at least 49 whole genome sequences, combining 1,092 strains in total (Table S1): 675 non-closely-related *S. cerevisiae* isolates from the 1,011 collection, 280 North American *S. paradoxus*, 88 globally distributed *S. eubayanus*, and 49 *S. uvarum* strains (Peter et al. 2018; Eberlein et al. 2019; Almeida et al. 2014; Nespolo et al. 2020). Mutations were polarized using either an outgroup species or an early-diverging population within each species (Materials and Methods). To ensure consistency of variant filtering across species, we reprocessed all raw sequencing reads (Table S1), called variants with a unified pipeline, normalized mutation frequencies by GC content of SNP sites used for mutation spectrum for each species, and performed PCA on these normalized spectra (Figure 1A). Notably, the mutation spectrum divergence between the African Beer and non-African Beer *S. cerevisiae* populations (adjusted variance = 0.001953) exceeds that observed between *S. eubayanus* and *S. paradoxus* (8.8690e-06; Materials and Methods). Similar results are obtained when performing the permutation-based AMSD method (Hart et al. 2026): cosine similarity is 0.0205 (p<1e-4) between African Beer and non-African Beer *S. cerevisiae* populations, while 0.00037 (p<1e-4) between *S. eubayanus* and *S. paradoxus.* Strikingly, the first principal component of *Saccharomyces* mutation spectrum variation separates the African Beer strains from a cluster that contains all non-*S. cerevisiae* species as well as all other *S. cerevisiae* strains. This deviates from the usual tendency for phylogenetically related lineages to have similar mutation spectra relative to more distantly related lineages, a pattern observed among mammalian species (Beichman et al. 2023; Goldberg and Harris 2022; Talenti et al. 2025).

To evaluate the effects of *de novo* variant calling and GC normalization, we compared these results with mutational PCA analyses performed using pre-filtered VCFs (Supplementary Figure 1) or generated with our unified pipeline without GC normalization (Supplementary Figure 2). In all three PCAs, the African Beer population consistently appeared as an outlier, and the relative positioning of the species’ spectra was broadly similar. However, we observed several minor discrepancies. Notably, mutational spectra from different species clustered much more closely after GC normalization. The stronger species separation observed in the pre-filtered or unnormalized analyses may therefore reflect technical artifacts, underscoring the importance of these normalization steps for cross-species comparisons.

African Beer strain variation appears enriched for A>G mutations along PC1 and for A>C mutations along PC2 (Figure 1A). To qualitatively estimate the timing of this enrichment, we examined how the mutation spectrum varies as a function of derived allele frequency and quantified the fold differences (odds ratio) between African Beer and French Dairy relative to the average spectrum of other *S. cerevisiae* populations (Figure 1B-C, Supplemental Figure 3). As higher derived allele frequencies generally indicate older alleles, this analysis provides insight into how long ago the African Beer mutation spectrum diverged from that of other *S. cerevisiae* strains. In African Beer, A>C mutations show a pronounced enrichment at intermediate allele frequencies near 0.01, peaking at nearly two-fold, followed by a decline at lower frequencies. However, enrichment remains elevated (∼1.5-fold) in the lowest-frequency bin relative to higher frequencies, suggesting that the mutational bias may still be active and affecting newly arising variants. This contrasts with a transient “pulse” as seen in the human TCC>TTC triplet mutation where the lowest-frequency bin no longer shows enrichment (Harris and Pritchard 2017). A>G, on the other hand, shows a transient and more modest enrichment, with only marginal enrichment within the lowest-frequency bin (Figure 1B-C). In French Dairy, the more recent allele frequency bins are also enriched for A>C and A>G, though the signal is weaker than it is in the African Beer population (Figure 1B-C). In addition, the hump from A>C and A>G in French Dairy is shifted toward lower frequencies than the broad hump from that of African Beer (Figure 1B, Supplemental Figure 3). Note that the absolute frequency of A>C is much lower than A>G, and their relative frequency is highest in African Beer, followed by French Dairy and other *S. cerevisiae* populations (Figure 1C), a pattern consistent across allele frequency bins.

To identify mutation signatures unique to the African Beer and/or French Dairy populations, we decomposed mutation frequencies into mutational signatures using SigProfilerExtractor (Islam et al. 2022) from the 1011 *S. cerevisiae* dataset, obtaining mutational triplet context from the *S. cerevisiae* reference genome. The African Beer population exhibited the most pronounced divergence in the decomposed mutational signatures. (Supplementary Figure 4). Strikingly, whereas most strains’ spectra were dominated by four major signatures—SBS1, SBS5, SBS12, and SBS102—more than half of the African Beer strains exhibited only three detectable signatures: SBS1, SBS5, and SBS12. In strains that are depleted of SBS102, some also showed nearly double the proportion of SBS5 relative to other *S. cerevisiae* strains, further distinguishing the African Beer population from all other strains.

Overall, the distinctive mutation spectrum of the African Beer strain appears to be a longstanding feature of its population of origin (Figure 1C, Supplemental Figure 3). In contrast, this signature seems to have appeared relatively recently in the French Dairy population, as evidenced by the rare allele frequencies of the excess A>G and A>C mutations, as well as the fact that the signal is only present in a subset of the French Dairy strains from the PCA analysis. We hypothesized that this pattern could reflect recent admixture from the African Beer population into the French Dairy population. In contrast, extreme divergence of the African Beer mutation spectrum is also likely to reflect either older introgression from an unknown “ghost” lineage, the long-term segregation of mutator alleles within this population, or a longstanding environmental mutagen. To distinguish among these possibilities, we performed additional analyses of the allele sharing between the French Dairy and African Beer strains.

### Admixture can produce intermediate mutation spectra on a mutational PCA

To illustrate that admixture can lead to an intermediate mutation spectrum, we examined strains from North American wild populations of *S. paradoxus*, which have relatively simple and well-characterized population histories (Eberlein et al. 2019). The SpD lineage arose from a recent hybridization event between the SpB and SpC* populations and occupies an intermediate position between them in the mutation spectrum PCA (Figure 2A). Because this hybridization event is recent, estimated to occur roughly 2600 years ago, the SpD lineage has had limited time to accumulate new mutations, suggesting that its intermediate mutation spectrum primarily reflects admixture rather than novel mutational processes.

**Figure 2.**
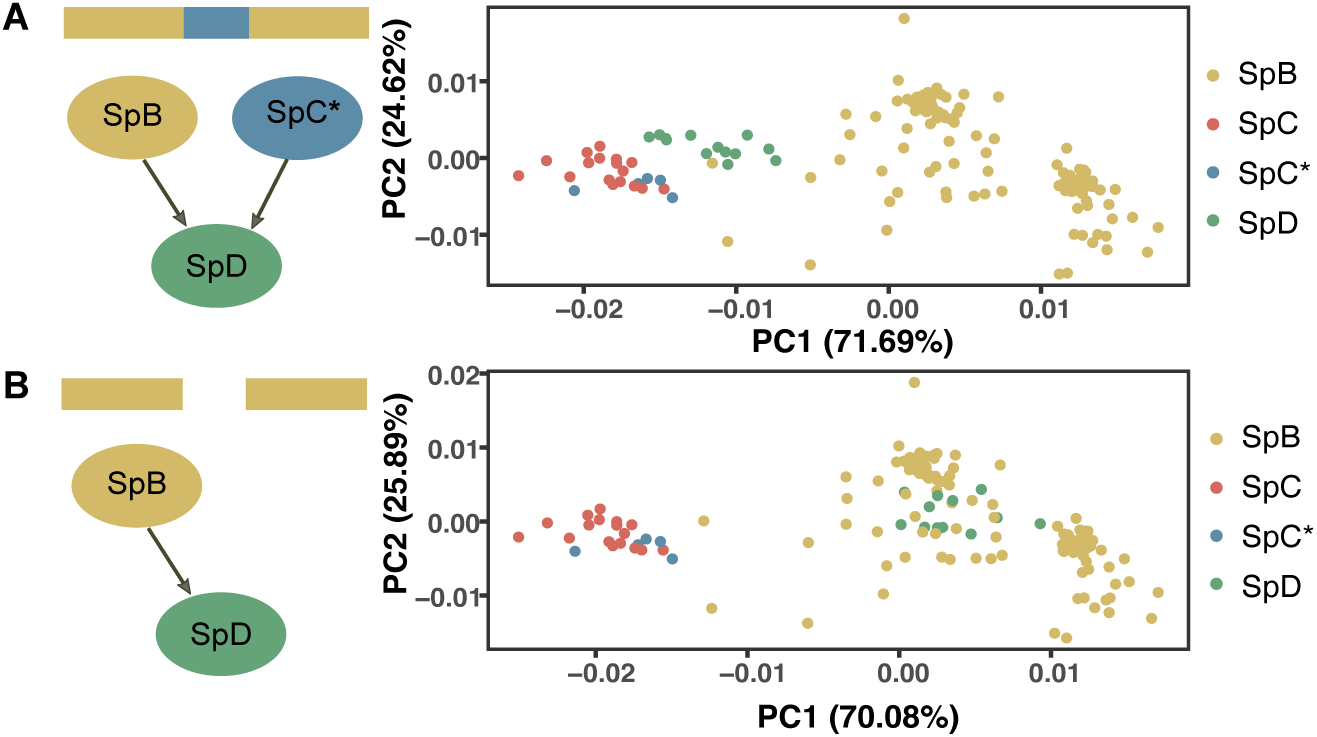
Admixture can cause intermediate positioning on the mutational PCA. (A) Mutational PCA from SNPs of North American *S. paradoxus,* polarized by SpA as the outgroup. SpD population shows an intermediate mutation spectrum compared with the two donor populations SpB and SpC*. (B) Mutational PCA of SNPs of North American *S. paradoxus,* except for SpD individuals, where only SNPs in the genomic region of SpB ancestry are counted. SpD population shows a similar mutation spectrum as SpB after removing SpC* ancestral genomic regions.

To demonstrate how admixture can explain mutation spectrum variation in *S. paradoxus*, we used Chromopainter2 (Lawson et al. 2012) to infer the ancestry of genomic tracts in SpD individuals, assigning each region to either the SpB or SpC* population. Our reconstructed ancestry patterns (Supplementary Figure 5) closely matched those inferred by Eberlein et al. (2019) using alternative methods. If the mutation spectrum of SpD can be explained as a simple function of its origin as an admixed population, then removing genomic regions inherited from SpC* should cause the remaining (SpB-derived) portions of the SpD genome to resemble the SpB mutation spectrum. To test this hypothesis, we recalculated the SpD mutation spectra using only the SpB-assigned genomic tracts from SpD individuals (Figure 2B). As predicted, the resulting spectra closely matched those of the SpB population (Green dots overlap with yellow), demonstrating that admixture alone can explain the relationship among the SpB, SpC*, and SpD mutation spectra in PCA space.

### French Dairy strain mutation spectra are inconsistent with a simple model of admixture from African Beer yeast

Since admixture can cause a hybrid’s mutation spectrum to appear intermediate between those of its founder populations, we revisited the 1011 *S. cerevisiae* dataset to investigate whether such a mechanism can explain why a subset of French Dairy mutation spectra appear intermediate between those of the African Beer population and other *S. cerevisiae* isolates (Figure 3A-B). We designated the subset of French Dairy strains that resemble the majority of other *S. cerevisiae* as French Dairy 1 (Figure 3D, yellow), and the subset with intermediate mutation spectra as French Dairy 2 (the 4 strains AQM, BFN, ARN, and BME shown green in Figure 3D). We found that French Dairy 2 exhibited an intermediate position among French Dairy 1 and African Beer using genome-wide SNP markers (Supplementary Figure 6), as expected under an introgression scenario.

**Figure 3.**
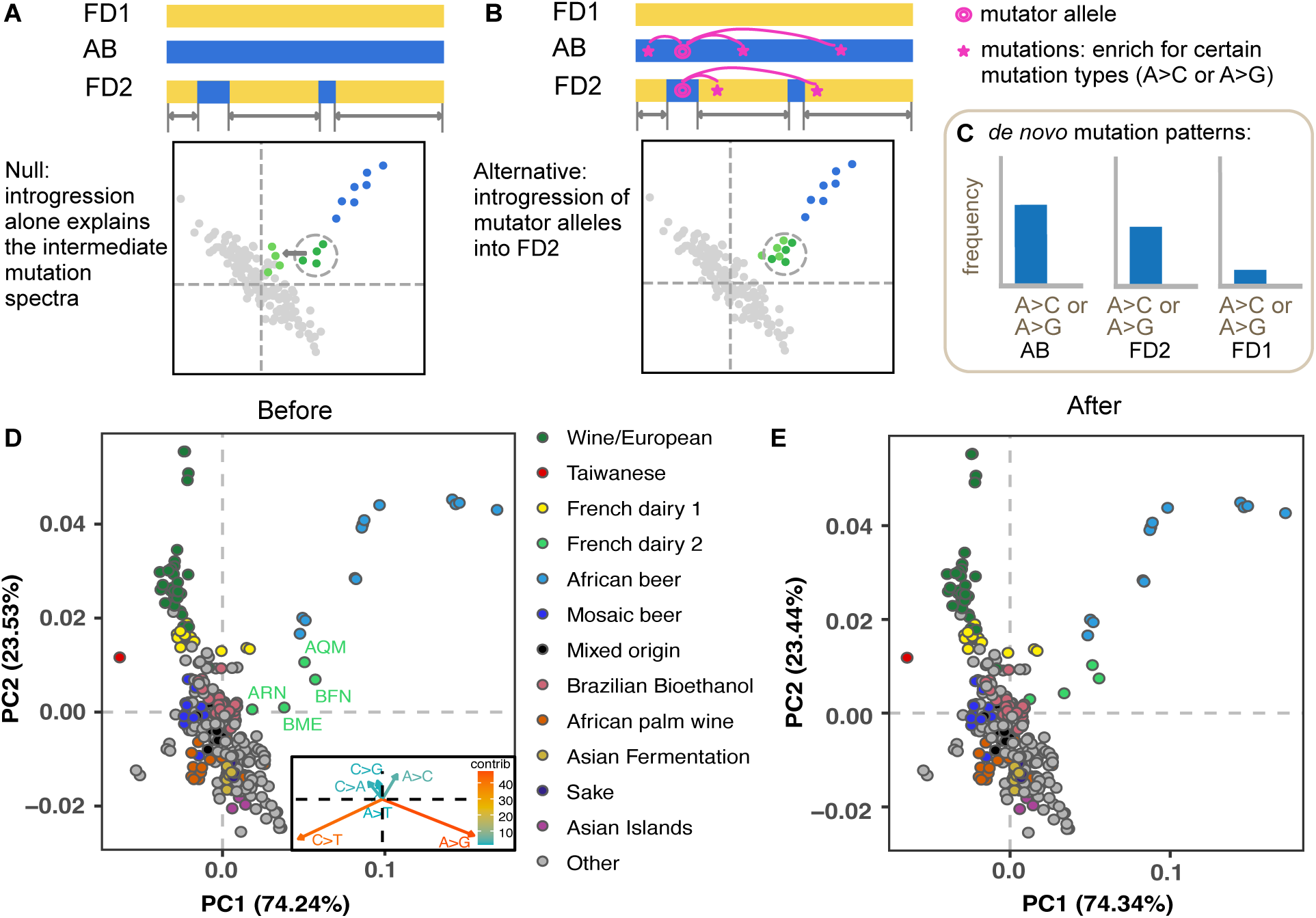
Removal of African ancestry tracks does not shift French Dairy 2 mutation patterns to overlap French Dairy 1. (A) Schematic diagram that predicts the behavior of French Dairy 2 strains on mutational PCA when the intermediate mutation spectra are solely due to admixture of African Beer DNA after removing African Beer ancestry tracks. (B) Schematic diagram that shows the predicted outcomes when there is an active mutator allele in African Beer DNA that got introgressed into French Dairy 2 strains. (C) Prediction of *de novo* mutation patterns according to the introgression of the mutator hypothesis in French Dairy 2. (D) Mutation spectrum PCA from 1011 *S. cerevisiae* strains, as in (Jiang et al. 2021), with the French Dairy group subdivided into French Dairy 1 (yellow) and 2 (green). (E) Mutation spectrum PCA from 1011 *S. cerevisiae* strains, after removing predicted African Beer ancestry tracks in the French Dairy 2 strains.

To more directly test whether the French Dairy 2 strains are products of admixture of an African- Beer-like yeast into the French Dairy population, we applied the D-statistic test for gene flow using the qpDstat tool from AdmixTools (Patterson et al. 2012). The four-population test was configured as (W, X, Y, Z) = (French Dairy 1, French Dairy 2, African Beer, outgroup), with outgroups from either the Mosaic Beer or African Palm Wine populations. Negative D-statistic values supported gene flow from African Beer into French Dairy 2 (*D* = -0.1202, *Z* = -6.426 using Mosaic Beer as outgroup; *D* = -0.2347, *Z* = -52.355 using African Palm Wine as outgroup).

After establishing that the French Dairy 2 strains are the product of admixture between African Beer and French Dairy 1 populations, we used Chromopainter2 to infer which segments of the French Dairy 2 genomes appear to originate within each ancestral population (Supplementary Figure 7). We then recalculated the French Dairy 2 strains’ mutation spectra after removing the putative tracts of African-Beer-like ancestry (Figure 3A-B). If the intermediate mutation spectrum of French Dairy 2 is solely due to admixture, removing the African Beer admixture tracts should render the spectrum similar to that of the French Dairy 1 strains (Figure 3A). Unexpectedly, removing African Beer tracts had little impact on the mutation spectra of French Dairy 2 strains, which retained their affinity with the African Beer strains after exclusion of all putative African Beer admixture tracts identified by Chromopainter2 (Figure 3D). This implies that the French

### Dairy 2 strains have African-Beer-like mutations that occur in DNA regions that are not readily identifiable as admixed from an African-Beer-like population

One possibility to explain such a pattern is that our Chromopainter2 analysis failed to identify major African Beer ancestry tracts, and that these unidentified tracts are responsible for the African-Beer-like mutation spectrum bias of the redacted French Dairy 2 genomes. We therefore repeated the D-statistic test on the redacted genomes and found that the D values moved closer to zero after ancestry tract redaction. When we used Mosaic Beer yeast as the outgroup, the *D*- statistic value changed from (*D* = -0.1202, *Z* = -6.426) to (*D* = -0.0863, *Z* = -4.673). Using African Palm Wine yeast as the outgroup, the *D*-statistic changed from (*D* = -0.2347, *Z* = - 52.355) to (*D* = -0.1365, *Z* = -12.332). These results indicate that removal of the Chromopainter2 tracts reduced the contribution of DNA related to the African Beer yeast by about 30 to 50 percent. However, our PCA analysis suggests that redaction reduced the African Beer mutational signature to a much lesser extent, indicating that this mutational process must also affect DNA that is not derived from recent admixture.

Given the above evidence, we hypothesize that the presence of the African Beer mutational signature throughout the French Dairy 2 genomes is due to introgression of at least one mutator allele from the African Beer population into the French Dairy population (Figure 3B). Such a mutator allele could influence the spectrum of *de novo* mutations arising in admixed genomes, causing the African Beer mutational signature to arise in French Dairy yeast. If this hypothesis is correct, then the spectrum of mutations originating *de novo* within the French Dairy 2 population should exhibit enrichment of the same mutation type(s) as African Beer (i.e., A>C and/or A>G, Figure 3C). In contrast, if the African Beer mutational signature is caused by an environmental mutagen or the transmission and subsequent loss of a mutator allele, we would expect the French Dairy 2 *de novo* mutation spectrum to resemble the French Dairy 1 mutation spectrum rather than the intermediate spectrum of the mutations that segregate within this population.

### Genomic engineering and examination of *de novo* mutations in African Beer and French Dairy strains

To test whether the African Beer and French Dairy 2 populations share mutational biases consistent with active mutator alleles, we analyzed the spectra of *de novo* mutations in natural isolates from three groups: African Beer, French Dairy 1, and French Dairy 2. Specifically, we aimed to experimentally compare the frequencies of A>C and A>G mutations across these populations.

We measured mutation spectra using a modified fluctuation assay based on *CAN1* reporter selection (Jiang et al. 2021). Because *CAN1*-based assays require a single functional copy of the sensitive allele and are typically applied to haploids, we engineered the diploid French Dairy and polyploid African Beer strains by deleting the endogenous *CAN1* loci with CRISPR and reintroducing a single sensitive *CAN1* copy tightly linked to a *NatMX* marker with transformation (Figure 4A; Table S2). This dual selection design avoids the detection of mitotic recombination events, which occur at much higher frequencies (∼10^-5^) (Thornton and Johnston 1971; Lee et al. 2009) compared with true *de novo* mutations (∼10^-7^) (Lang and Murray 2008). We successfully constructed all reporter strains, confirming through PCR and sequencing that each carried a single *CAN1–NatMx* copy at the target locus. We then collected and sequenced pooled mutants following growth in MSG/SC-Arg + 2% glucose media, selection and regrowth on canavanine + clonNAT, and PCR amplification of the *CAN1-NatMx* locus of pooled mutants (Figure 4B; Materials and Methods). Since some of the African Beer strains are flocculant, reliably estimating their mutation rates was difficult, but we still expect our assay to yield unbiased measurements of their mutation spectra.

**Figure 4.**
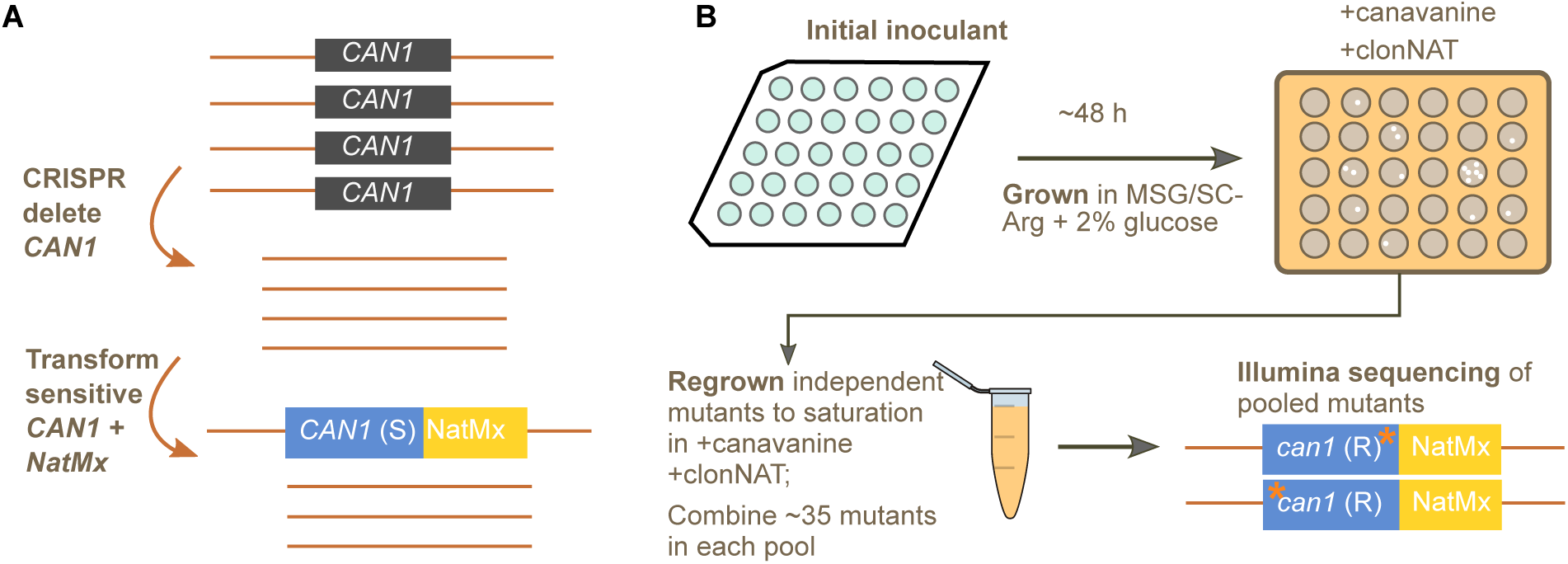
**Experimental pipeline to measure *de novo* mutation spectra from African Beer and French Dairy strains**. (A) Genetic engineering of African Beer and French Dairy strains to introduce one sensitive copy of the *CAN1-NatMx* reporter. This shows an example of a tetraploid strain, such as ANM. (B) Modified fluctuation assay that captures the mutation spectrum of each strain with the selection of both *can1* and *NatMx*.

As a control, we engineered a haploid laboratory strain LCTL1 (*SEY6211-MATα*) along with French Dairy and African Beer strains (Table S2) following the same procedure, which was previously characterized alongside experimental strains using the original version of our fluctuation assay method (Jiang et al. 2021). To ensure we recapitulate the same mutation spectrum of the unmodified strain from the previous work, we examined ∼300 mutants from the control strain LCTL1. This revealed a mutation spectrum that differed slightly but significantly from that of the unmodified strain (*p* = 0.023; Supplementary Figure 8A). This discrepancy could reflect either biases introduced during regrowth of mutant colonies under *NatMx* selection, or direct effects of the modified medium affecting the mutation process itself. To distinguish between these possibilities, we repeated the assay by replacing with canavanine-only media during the regrowth of canavanine-resistant mutants. Under these conditions, LCTL1 showed near-complete regrowth (∼100%) compared to ∼50% in canavanine + clonNAT, while natural isolates tested (French Dairy or African Beer strains) were unaffected with a near-complete regrowth rate in canavanine + clonNAT (Supplementary Figure 8B). The mutation spectrum of LCTL1 in canavanine-only conditions closely matched that of the unmodified *CAN1* locus (n.s., Supplementary Figure 8A), indicating that the altered spectrum arises from differential regrowth under dual selection rather than changes in mutagenesis. This effect appears specific to the LCTL1 background, likely reflecting epistatic interactions that reduce growth of certain mutants under canavanine + clonNAT. Importantly, no such regrowth bias was observed in French Dairy or African Beer strains, indicating that their mutation spectra are not confounded by this artifact.

### *de novo* mutation of A>C type is consistent with the mutator allele introgression from African Beer into French Dairy 2

We collected approximately 300 mutants of each engineered natural isolate, identified mutations (Table S3), and calculated their mutation spectra. Examination of the full spectra revealed that a haploid African Beer strain AFL was dominated by C>A mutations (∼80%) (Figure 5B). This pattern suggests that AFL carries an additional, strong mutator allele that is not characteristic of the African Beer population as a whole and specifically elevates the C>A mutation rate, thereby obscuring other mutational signals. Consequently, AFL was excluded from subsequent analyses.

**Figure 5.**
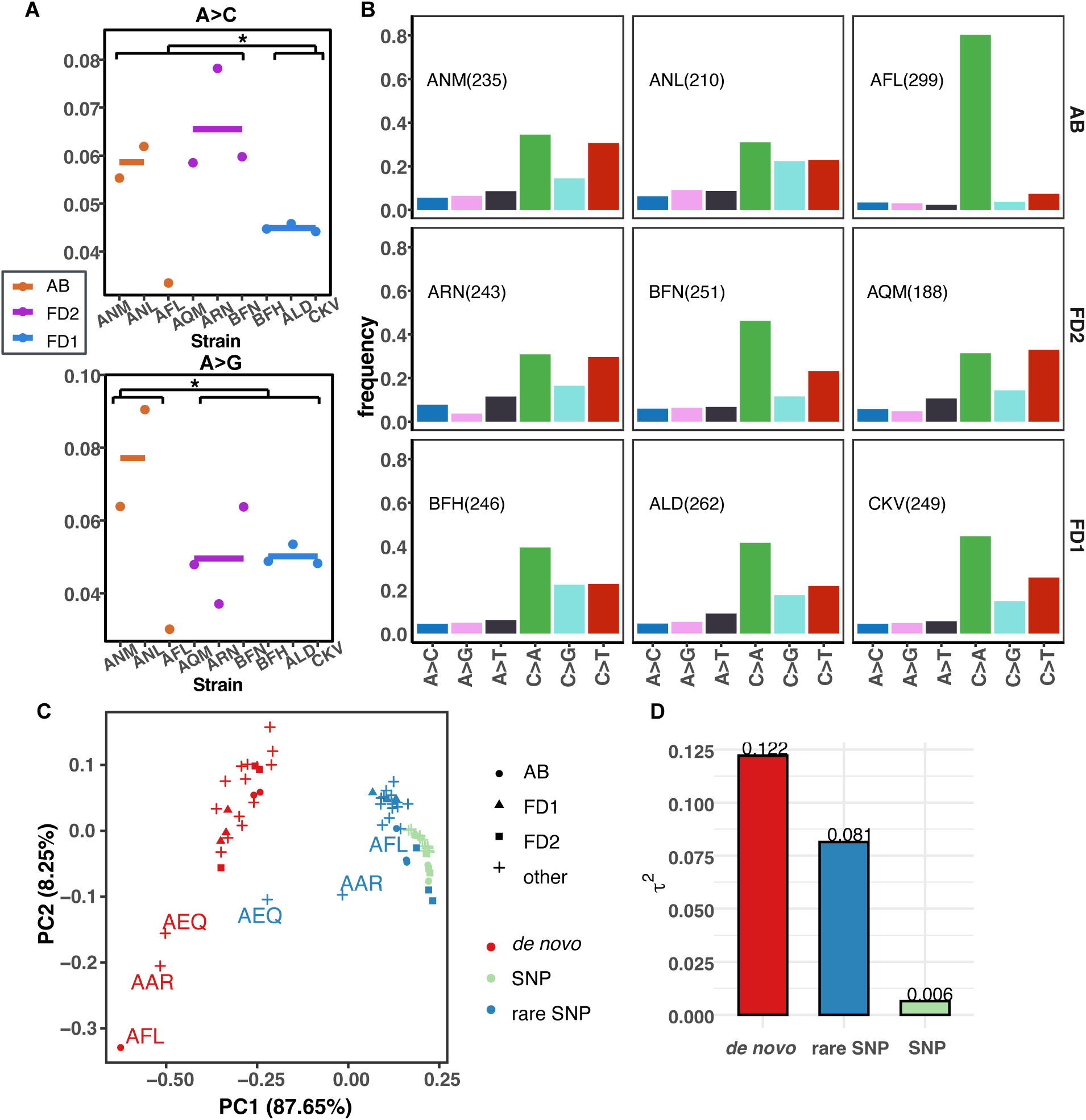
*de novo* mutation of A>C in African Beer and French Dairy strains is consistent with the prediction of mutator allele(s) in African Beer. (A) *de novo* mutation of A>C and A>G type in African Beer (AB), French Dairy 1 (FD1), and French Dairy 2 (FD2) populations. (B) *de novo* mutation spectrum of engineered African Beer, French Dairy 1, and French Dairy 2 strains. (C) PCA analysis of mutation spectrum for *de novo*, rare SNPs, and all SNPs in African Beer, French Dairy 1, and French Dairy 2. Strains that exhibit outlier mutation spectrum in the previous study: AEQ and AAR, and in this study: AFL, are labeled. (D) Comparison of Aitchison’s total variance *τ*^2^ of the mutation spectrum among *de novo*, rare SNPs, and all SNPs.

The fractions of A>Cs across the African Beer, French Dairy 1, and French Dairy 2 groups (Figure 5A) were broadly consistent with our expectations given the hypothetical introgression of a mutator allele from African Beer into French Dairy 2. On average, African Beer and French Dairy 2 strains showed roughly a 30% higher A>C mutation frequency than French Dairy 1. Using the CLR-based transformation of the mutation spectrum, followed by PERMANOVA (Anderson 2017), A>C is significantly different among the three groups (*p* = 0.0207). We also see a significant difference in the A>C mutation frequency between the French Dairy 1 and the supergroup containing African Beer and French Dairy 2 (*p*=0.0178). We further applied the AMSD test (Hart et al. 2026) on the combined dataset, and obtained the same results (*p*=0.0187). This pattern is consistent with the hypothesis that a mutator allele influencing A>C mutations was introgressed from African Beer into French Dairy 2 and remains active. In contrast, A>G mutation frequency differed significantly only between the African Beer population and the French Dairy groups (*p* = 0.0336), rather than showing the shared enrichment in African Beer and French Dairy 2 expected under the introgression hypothesis.

When examining the complete *de novo* mutation spectra, the differences in A>C and A>G ratios among strains were modest. In contrast, greater variability was observed across strains for other mutation types (Supplementary Figure 9, A>C and A>G have the lowest adjusted variance of 0.027 and 0.054 respectively, and the variance in other mutation types varies between 0.055 and 0.294). It is likely due to the fact that A>C and A>G are among the rarest mutation types. This suggests that while multiple mutator alleles may arise and affect mutation spectra, only those with mild effects, which have limited fitness consequences, are likely to persist. To test this hypothesis, we compared the variance in mutation spectra derived from *de novo* mutations, rare SNPs, and all SNPs across natural yeast isolates characterized in this and previous work via mutational PCA and variance analysis ((Jiang et al. 2021); Figure 5C). As predicted, Aitchison total variance was highest for *de novo* mutations, intermediate for rare SNPs, and lowest for all SNPs (Figure 5D). This pattern indicates that strong mutator alleles arise transiently but are purged by selection, whereas mild mutators—such as those influencing A>C and A>G mutations—can leave detectable evolutionary signatures in natural populations, as observed in the African Beer group.

Since the enrichment of A>C mutations is consistent with the introgression of active weak mutator(s) acting in the recipient French Dairy strains, we sought to identify putative mutator loci in these genomes. We first identified genomic regions introgressed from African Beer strains that were shared among the four French Dairy 2 isolates (Table S4), encompassing 26 regions across 10 chromosomes. Within these regions, we detected 42 protein-coding genes (Table S5). Among these, three candidate genes — *SSL2 (YIL143C), RAD27 (YKL113C), APN1 (YKL114C) —* overlapped with a curated set of 158 genes previously known to function in DNA replication and repair (Jiang et al. 2021). *SSL2* functions in transcription initiation and nucleotide excision repair (Feaver et al. 1993). *RAD27* encodes a nuclease involved in DNA replication and multiple DNA repair pathways (Kao et al. 2002; Ayyagari et al. 2003; Sommer et al. 2008; Sparks et al. 2012). *APN1* encodes the major apurinic/apyrimidinic (AP) endonuclease in the base excision repair pathway (Ma et al. 2008). *apn1*-deficient yeast exhibit elevated single-base substitutions, including a striking 59-fold increase in A>C transversions in the *SUP4*-o reporter assay (Kunz et al. 1994).

## Discussion

This study reveals a novel pattern consistent with the long-term persistence and transmission of mild mutator alleles affecting A>C mutations from African Beer into French Dairy in natural yeast populations. Mutator alleles that strongly elevate mutation rates are typically deleterious and thus are rarely thought to persist over evolutionary time. However, our findings suggest that weak mutators—those with small effect sizes—can remain in populations and shape mutation spectra over extended periods. Although *de novo* mutation analyses revealed that many mutation types vary among strains, only the mild A>C and A>G biases associated with the African Beer group show evidence of accumulation regarding their natural polymorphism patterns.

Despite recent advances in uncovering the variation in mutation rate and spectrum in natural populations across species, there are limited studies linking patterns of mutation rate variation to the transmission of genetic mutators in natural populations. Most studies have focused on describing static differences in mutation spectrum polymorphisms among species or populations (Goldberg and Harris 2022; Beichman et al. 2023; Talenti et al. 2025). Studies that inferred historical mutational changes from polymorphism data have identified transient enrichments of specific mutation types within particular populations, such as the TCC>TTC pulse (Harris and Pritchard 2017; Talenti et al. 2025). However, the underlying causes of these enrichments remain elusive, and there is no evidence that such mutational processes are still segregating in extant populations.

Studies based on *de novo* mutations have revealed substantial variation in mutation rates and spectra among individuals and populations (López-Cortegano et al. 2024; Wang et al. 2023), as well as rare cases in which mutator alleles could be associated with altered mutation patterns (Kaplanis et al. 2022; Zhang et al. 2025). Nevertheless, these studies provide limited insight into the evolutionary history and long-term transmission of mutator alleles within populations. In yeast, our previous work (Jiang et al. 2021) identified a recently arisen natural mutator allele that influences both *de novo* mutations and rare natural polymorphisms. In mice, (Sasani et al. 2022, 2024) identified mutator alleles associated not only with *de novo* mutation patterns but also with elevated levels of polymorphism mutations in wild populations carrying these alleles, suggesting that some mutator alleles can persist over evolutionary timescales. This study provides additional evidence that mutator alleles can persist in natural populations for extended periods of time. Unlike previous studies that primarily identified transient mutational signatures or recently emerged mutators, our results suggest that weak mutator alleles may remain in the population while continuously shaping patterns of standing genetic variation.

Detecting weak mutators poses a particular challenge because their effects can be subtle relative to the stochasticity inherent to accumulating *de novo* mutation and can also be masked by stronger mutational signatures. In our experimental assays, each strain accumulated roughly 300 *de novo* mutations across three biological replicates, with A>C mutations representing less than 10% of the total mutation spectrum. While the effects we observed were modest, their direction was consistent across analyses: both African Beer and French Dairy 2 showed enrichment of A>C mutations from natural polymorphisms, supporting the hypothesis that weak mutator allele(s) were introgressed from African Beer into French Dairy 2. In contrast, the A>G enrichment appeared restricted to the African Beer population, suggesting either that the corresponding mutator allele(s) were not retained in French Dairy 2 or not introgressed into French Dairy 2. Alternatively, the African Beer-specific A>G enrichment may reflect environmental factors rather than an underlying genetic difference. These observations suggest that the determinants of A>C and A>G enrichment are most likely distinct and segregate independently. It is also possible that the African Beer population’s excess A>C and A>G have the same underlying cause with different magnitudes, but that our assay failed to reveal this due to limited resolution. The *de novo* A>C and A>G mutation counts are too small to be persuasive on its own, but these measurements add to the picture of mutation spectrum differentiation followed by introgression that is clearly revealed by our genetic variation analysis. On the other hand, experiments that focus on *de novo* mutations alone could pick up stronger signals, such as C>A enrichment in AFL, which may not contribute to the evolutionary history of the population.

Future studies are needed to validate putative mutator alleles. Among the three candidate genes we found, *SSL2* and *RAD27* are likely to have broad effects on genome stability, whereas *APN1* operates at a finer scale by repairing AP sites—common spontaneous lesions produced by DNA base loss or damage arising from intrinsic DNA instability and the cellular microenvironment (Auerbach et al. 2005; Kunz et al. 1994). The concordance between this elevated A>C transversion rate in *apn1Δ* mutants and the A>C enrichment observed in the African Beer and French Dairy 2 populations suggests that *APN1* could represent a natural weak mutator allele underlying the long-term mutational signature observed in these groups. It is also possible that additional genes from the introgressed regions, but are not annotated as DNA repair contributes to this pattern. Although weak mutator effects are very challenging to characterize experimentally and have less of a short term effect on fitness than classical mutator phenotypes, they are more likely than classic strong mutator alleles to persist over evolutionary time and shape broad patterns of natural genetic variation.

## Acknowledgement

PJ was supported by a Burroughs Wellcome Fund Career Award at the Scientific Interface awarded to KH. KH acknowledges additional support from a Searle Scholarship, a Sloan Research Fellowship, a Pew Biomedical Scholarship, and National Institute of General Medical Sciences Grant 1R35GM133428-01. The research of MJD was supported by NIH/NIGMS award R01 GM147040. MJD holds the William H. Gates III Endowed Chair in Biomedical Sciences.

## Materials and Methods

### Mutation spectrum analysis in *Saccharomyces*

Raw reads were downloaded from NCBI and aligned to the reference genome of each species using bwa mem (v. 0.7.17), followed by variant calling with GATK HaplotypeCaller and GenotypeGVCFs (v. 4.2.6.1), and samtools (v. 1.16). Variants with QD < 2.0, QUAL < 30, SOR > 3.0, FS > 60.0, MQ < 40.0, MQRankSum < -12.5, ReadPosRankSum < -8.0, DP<10 were filtered out (see snakemake pipeline on Github). Bi-allelic SNPs are included for mutation spectrum analysis. After obtaining mutation counts for the six. Mutation PCA is performed the same way as in (Jiang et al. 2021) using all single-bp variants with derived allele frequency <0.5 per strain. For the outgroup used for polarizing mutations from SNPs: for *S. cerevisae* populations, 5 strains of *S. paradoxus* were used (Jiang et al. 2021). For *S. paradoxus*, a strain LL2012_001A from the SpA population was used as the outgroup. For *S. eubayanes,* the reference sequence from *S. uvarum* was used as the outgroup. For *S. uvarum*, ZP_962, ZP_963, ZP_964, and ZP_966 were used as the outgroup. The GC are the proportions of GC in the SNPs in the reference genome that are used to calculate mutation spectrum for each species are used for normalization. Mutation frequency of p(A>C), p(A>G), p(A>T), p(C>A), p(C>T), p(C>G) are normalized such that, S= p(A>C)/(1-GC) + p(A>G)/(1-GC) + p(A>T)/(1-GC) + p(C>A)/GC + p(C>T)/GC + p(C>G)/GC, into the adjusted frequency p(A>C)/(1-GC)/S, p(A>G)/(1-GC)/S, p(A>T)/(1-GC)/S, p(C>A)/GC/S, p(C>T)/GC/S, p(C>G)/GC/S. For calculating the adjusted variance, the centered log-ratio (clr) transformation (Aitchison 1986) was performed using “compositions” package in R (van den Boogaart and Tolosana-Delgado 2008) for each GC-adjusted mutation frequency, and then Aitchison’s total variance *τ*^2^ per group was calculated. For the between-group adjusted variance, we used the between-group sum of squares after the clr transformation.

### Predicting ancestral tracks and mutation spectrum re-analysis

Chromopainter 2 (Lawson et al. 2012) is used to predict the most likely ancestral tracks for SpD in *S. paradoxus* and French Dairy in *S. cerevisiae.* For *S. paradoxus*, strains from SpB and SpC* were used as donors, and strains from SpD were the recipients. In each of the SpD strains, the mutations from the predicted SpC* ancestry were not included, and the mutation PCA was performed on all the strains with updated mutation counts. For *S. cerevisae*, African Beer and French Dairy 1 strains were used as donors, and French Dairy 2 strains were the recipients. In each of the French Dairy 2 strains, the mutations from the predicted African Beer ancestry were not included, and the mutation PCA was performed on all the strains with updated mutation counts. Unphased genotype data from the donor and recipient are used, and a uniform recombination rate is assumed.

### D-statistic analysis for Introgression

We include all strains in French Dairy 1, French Dairy 2, and African Beer populations as populations W, X, Y. We picked two outgroups based on the 1011 phylogeny – one from the Mosaic Beer clade, and the other from the African Palm Wine clade. The reason to pick these two groups is that they are the closest outgroups to French Dairy and African Beer when excluding the mostly admixed strains (Peter et al. 2018). qpDstat from Admixtools was used to calculate D-statistics for the 4-population test (Patterson et al. 2012).

### Strain construction

Table S1 lists all the strains engineered in this study. We first use CRISPR to delete all copies of endogenous *CAN1* in the genome. We PCR the 5’ and 3’ flanking sequence of *CAN1* using either French Dairy or African Beer backgrounds as the template to amplify. CRISPR plasmid and guide RNA design and cloning follow (Laughery et al. 2015). We designed gRNA for *CAN1* using online tools developed in Laughery et al, and included them if they are top hits in Benchling. Two guides were designed to delete endogenous *CAN1* and both work, most strains use guide 1586 to delete endogenous *CAN1*. (guide 1586: CAGGAAAATAGAGATATAGG, guide 99: ATGGTATTGACCCACGTCTG). We use pML104-NatMx as the CRISPR plasmid backbone (NatMx marker is added by Anja Ollodart in the Dunham lab from pML104). *CAN1* deletion template was constructed by amplifying 5’ and 3’ flanking sequences of *CAN1* of African Beer (BEK), French Dairy strain (BGR), and lab strain (GIL 104) (five-prime-forward primer: GGGTGACGTGAAGATAACGA, five-prime-reverse primer with overhang: GGCATAGCAATGACAAATTCGTTTTATTACCTTTGATCACATTTCC, three-prime-forward primer with overhang: GTGATCAAAGGTAATAAAACGAATTTGTCATTGCTATGCCTTT, three-prime-reverse primer: GGTTTCTGTGTGGTTTCCG), followed by assembly PCR with five-prime-forward primer and three-prime-reverse primer. Successful transformation of CRISPR plasmid by selecting for clonNAT resistance. Engineered *CAN1* deletion strains were confirmed through PCR and whole-genome sequencing.

We then PCR amplified the *CAN1*-NatMx from AH4003 (obtained from Dr. Alan Herr) and transformed it into the *CAN1*-deletion strains, with primers: CAN1-321F: GAACTCTTGTCCCTTATTAGCCTTGATAGTGC, CAN1R2: GGAGTTTCAAATGCTTCTACTCCGTCTGC for African Beer strains and primers cannat_953_for: CCGAGATACGATTACTCCAGTTCCTC, cannat_4764_rev: GGAGTTTCAAATGCTTCTACTCCGTCT for French Dairy strains, using Kapa HiFi Hotstart Ready Mix (Roche catalog number: KK2601). The *CAN1*-NatMx fragment is transformed into the *CAN1-*deleted strains. Successful transformation was confirmed with primers: can-int-out- rev-1: AGGGCCCATTATGAATACGCA, Nat-mid-for-1: GTCCGATTCGTCGTCCGATT.

### Fluctuation assay, sequencing, and mutation calling

Fluctuation assays were conducted the same way as (Jiang et al. 2021, 2022), with the following modifications: yeast cells were grown in MSG/SC-Arginine + 2% Glucose media (replacing 5g ammonium sulfate in SC with 1g monosodium glutamate compared with the equivalent SC-Arginine media) for 48 hours, and the selection plates contained MSG/SC-Arginine-Serine, plus canavanine (60 mg/liter L-canavanine) and 2x clonNAT (1x clonNAT: 100ug/ml). Individual colonies from different replicates were picked and regrown in MSG/SC-Arginine-Serine, with canavanine and 1x clonNAT media, followed by the same pooling and Illumina sequencing as (Jiang et al. 2021, 2022), by PCR the *CAN1-Nat* amplicon with primers: CAN1_CN_for1: TGGGCACAAACCCTTGGAAA, CAN1_CN_rev1: TTCCAGCCGGAGTACGAGA with Kapa HiFi Hotstart Ready Mix. The alternative regrowing canavanine-only media contains MSG/SC-Arginine-Serine, plus canavanine.

The mutation calling pipeline is similar to (Jiang et al. 2021, 2022) with the following modifications: 1) regions 974-1000 of *CAN1-NatMx* amplicon were excluded due to putative false positive signals for the mutation spectrum for the *CAN1-Nat* amplicon. 2) Multi-nucleotide mutations (MNMs) were not excluded when calculating for mutation spectrum because they reflect the mutations that are accumulated. Raw reads of the pooled sequencing will be available at NCBI SRA upon publication. The full data analysis pipeline is available on GitHub (https://github.com/pyjiang/french_dairy).

