## Supplemental Figure 1-9 for "Evidence for long-standing mild-effect mutator alleles shaping genetic variation in *S. cerevisiae* populations"

Supplementary Figure 1

Mutation spectrum PCA when using pre-filtered VCF files for each species.

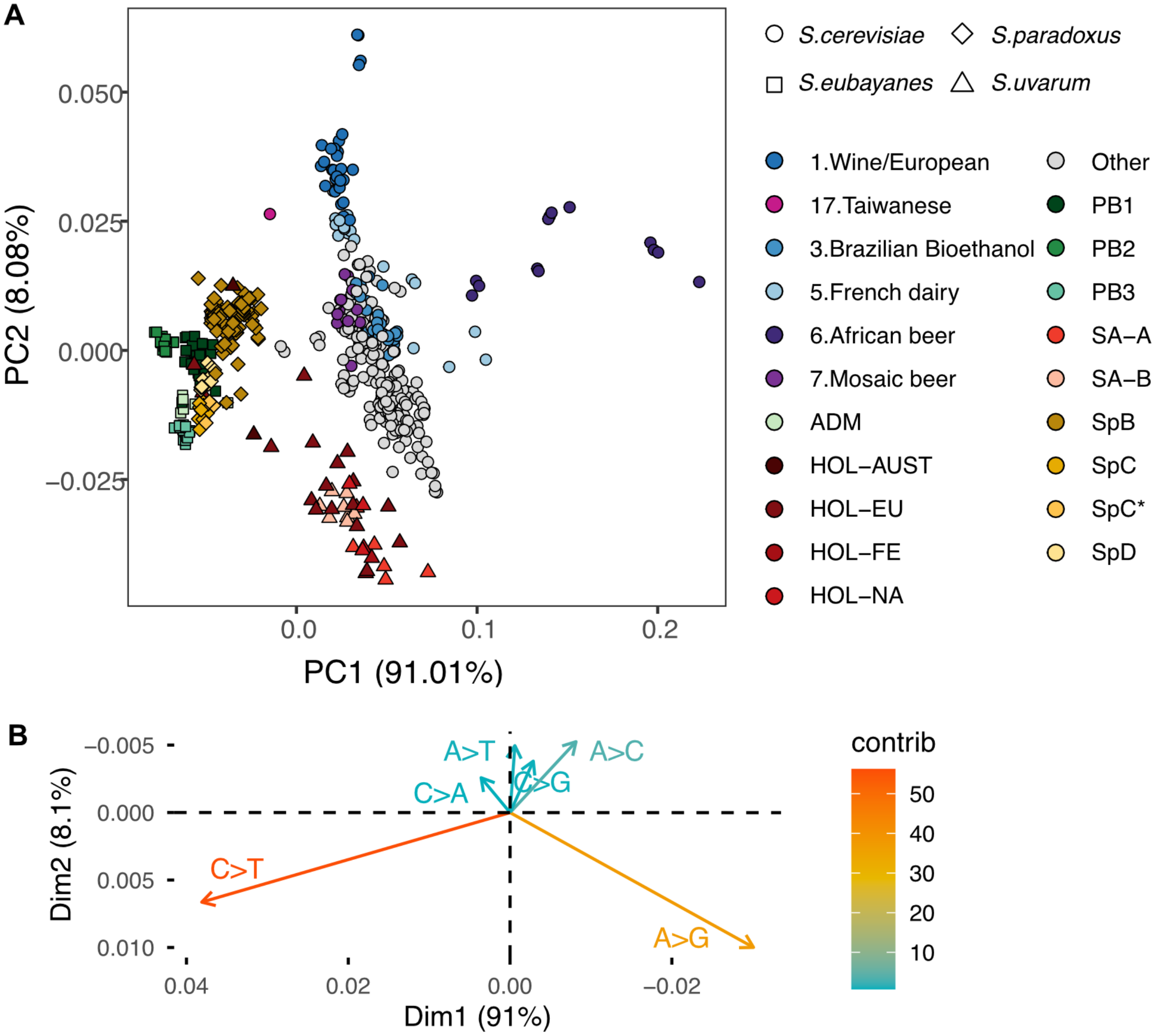

Supplementary Figure 2

Mutation spectrum PCA using variants called from raw reads with a common pipeline, without GC normalization.

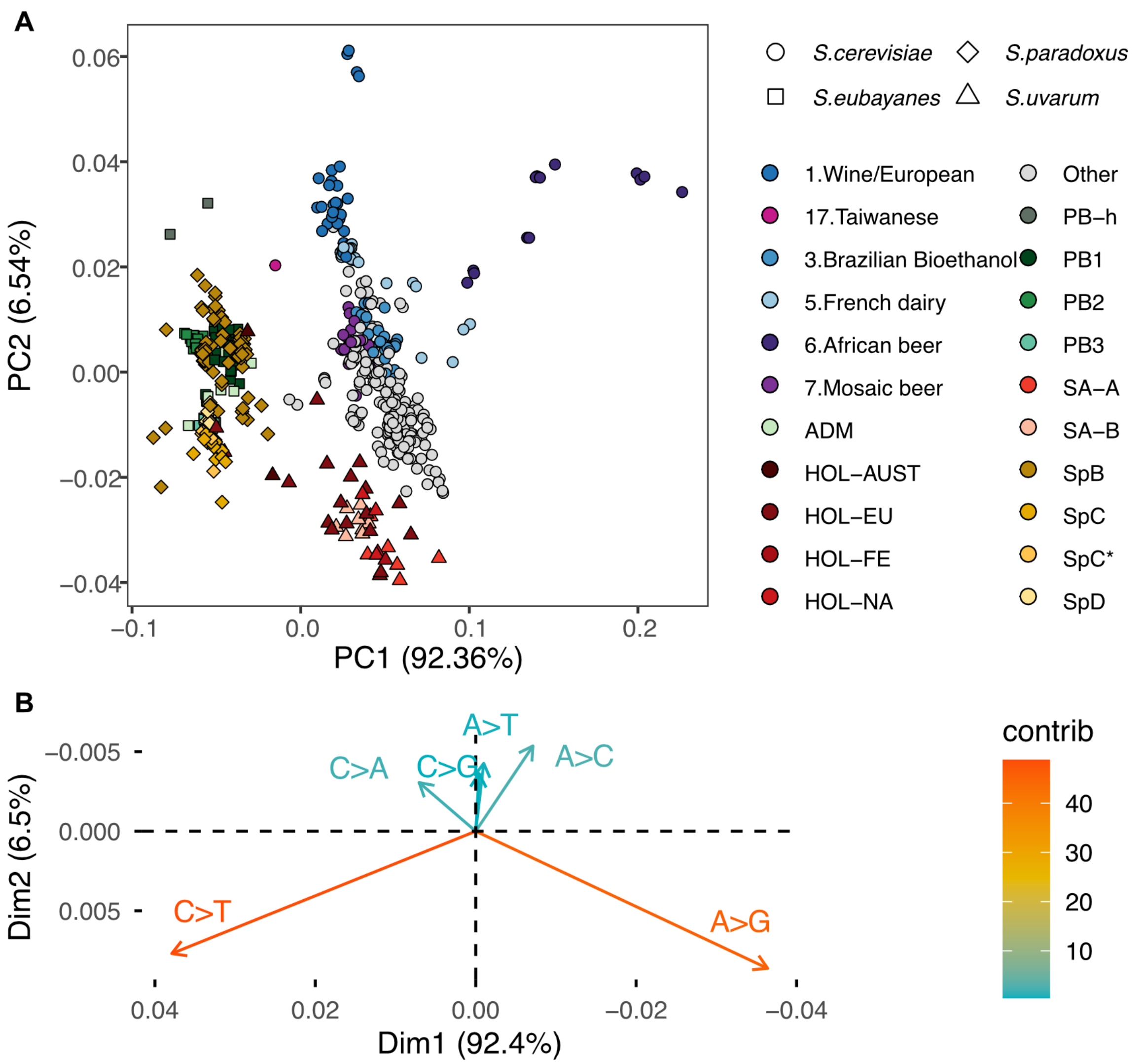

Supplementary Figure 3

Comparison of the mutation frequency of African Beer, French Dairy, and all other S. cerevisiae among derived allele frequencies.

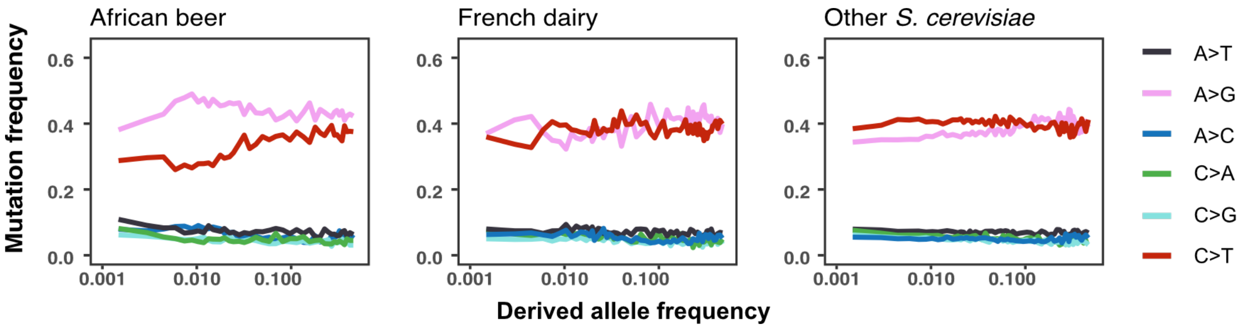


Supplementary Figure 4

Mutation signature decomposition from polymorphisms of *S. cerevisiae* strains.

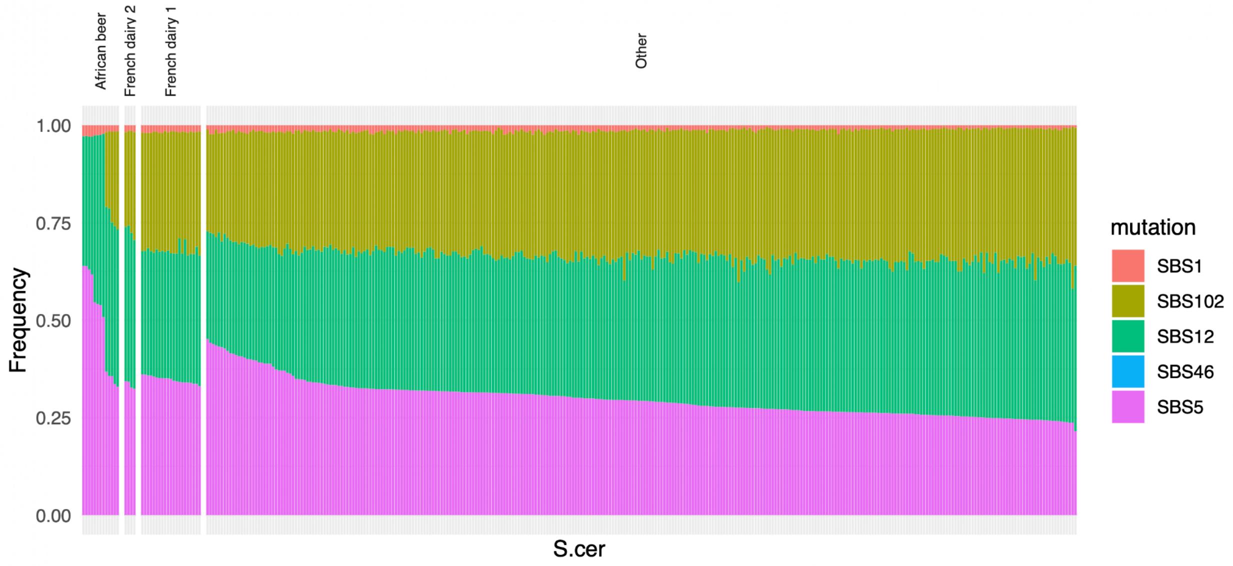

Supplementary Figure 5

Predicted tracks for SpD individuals from donors (SpC*: blue, SpB: yellow) in *S. paradoxus.*

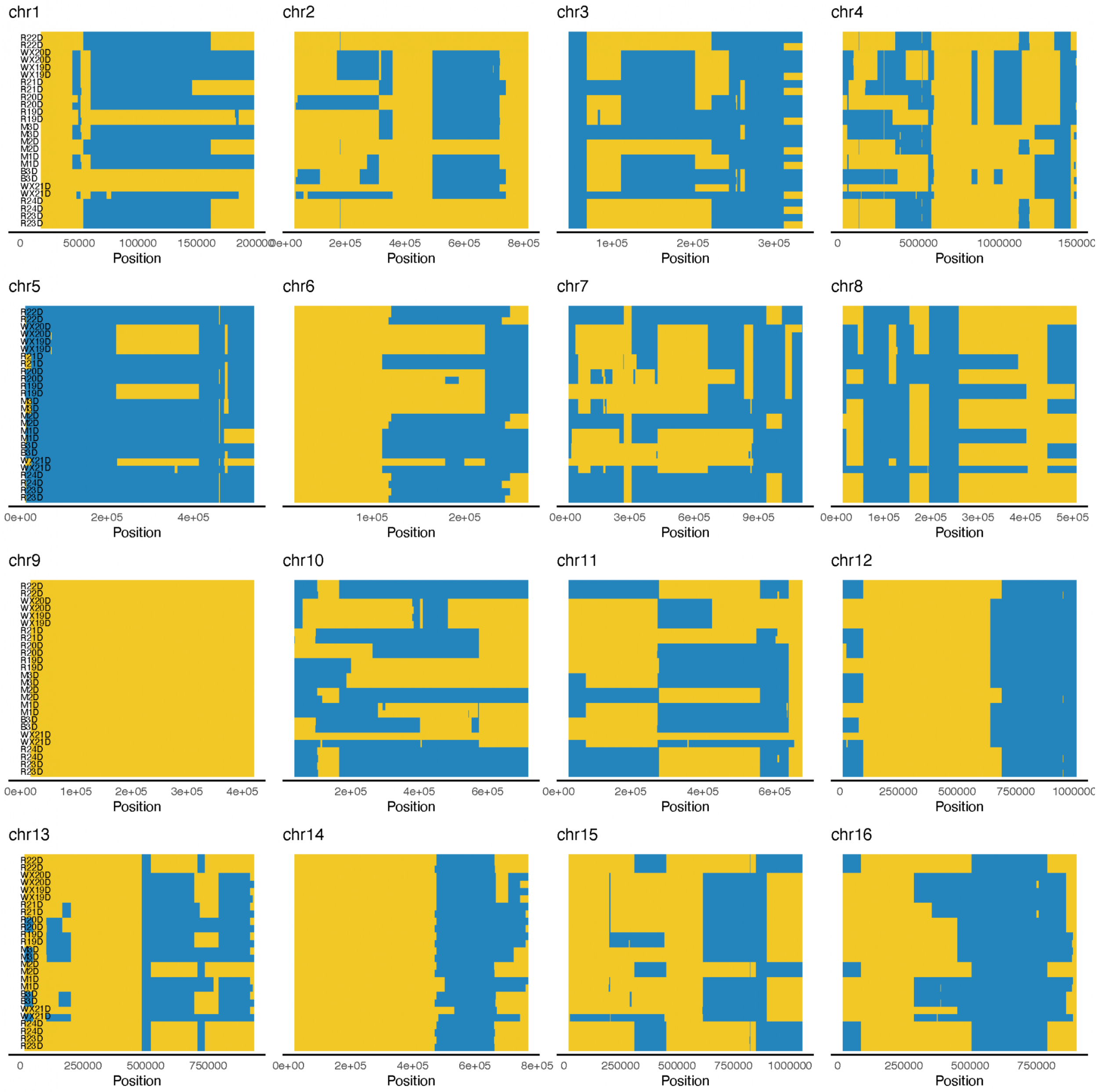

Supplementary Figure 6

PCA of SNPs in African beer and French dairy populations.

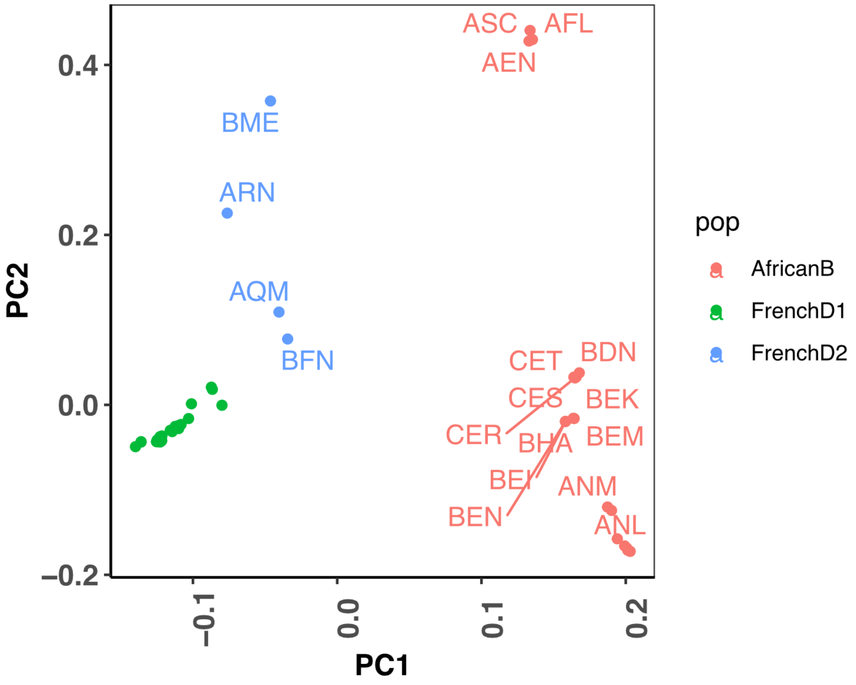

Supplementary Figure 7

Predicted tracks for French dairy 2 individuals from donors (African beer: blue, French dairy 1: yellow).

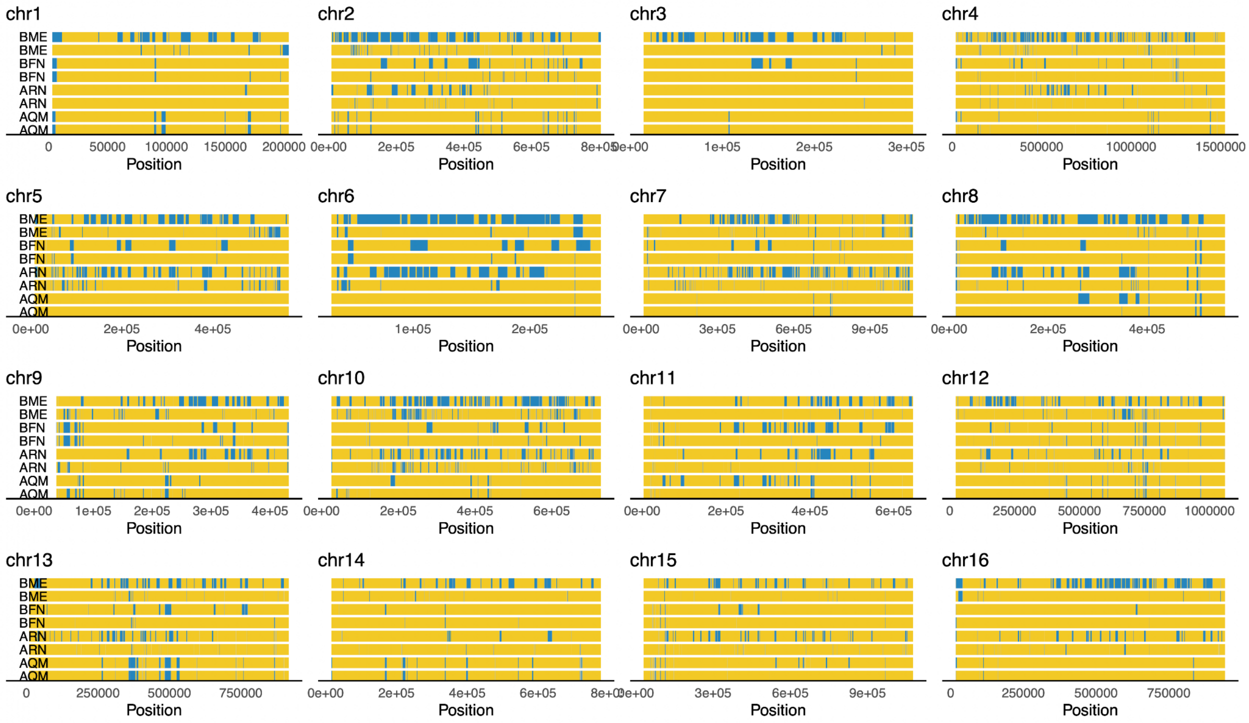


Supplementary Figure 8

Explanation of the discrepancy in the mutation spectrum of the control strain

(A) Mutation spectrum of control strain LCTL1 and of mutants that regrow in canavanine+clonNAT media or canavanine-only media, compared to the unmodified LCTL1. (B) Percentage of mutants that regrow in canavanine plus clonNAT media or canavanine-only media among testing strains and the control lab strain LCTL1. Two replicates are shown for LCTL1.

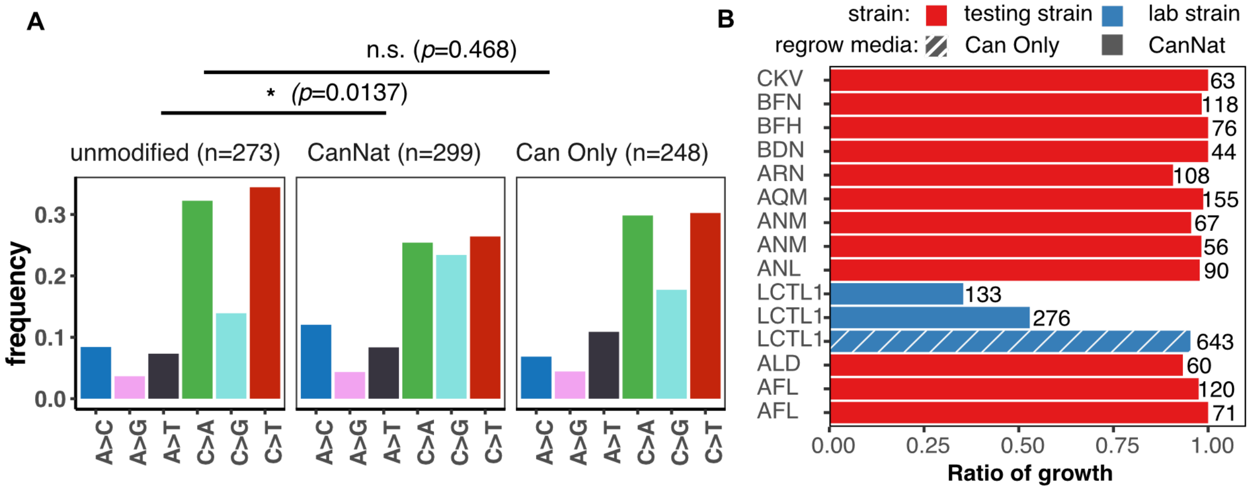

Supplementary Figure 9

Adjusted variance (variance after CLR transformation of mutation frequency) for different mutation types across all *de novo* mutations in this study

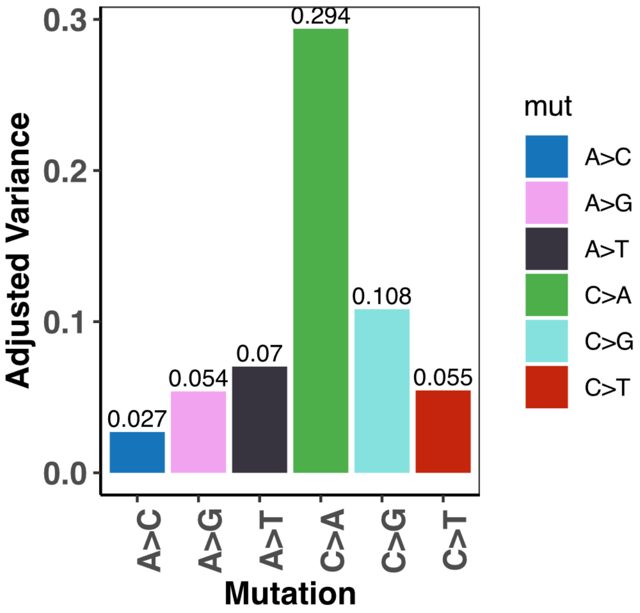
